# Middle-income countries disproportionately face the burden of the Open Access publication model

**DOI:** 10.1101/2025.10.23.684097

**Authors:** Imran Samad, Divyashree Rana

## Abstract

While the Open Access (OA) publishing model may promote equity in the readership of scientific publications, its implications for global research authorship remain poorly quantified. We analysed 1,117,978 publications over the past 20 years from 587 ecology and evolution journals to compare differences in publication rates, a) between Subscription-Based (SB) and OA publishing models, and b) before and after journals transitioned to the fully OA model, across country income groups and journal Article Publishing Charges. We found that, compared to the SB model, publication rates were generally lower under the OA publishing model, and these differences were highest in middle-income countries (63%, where only APC discounts are provided) than in low-income countries (43%, where full APC waivers are automatically provided). These differences also increased with increasing journal APC. For journals transitioning to a fully OA publication model, publication rates decreased for journals with low APCs (<$2425 USD) but increased for journals with higher APCs (<$3190 USD). We argue that as OA impacts publishing potential based on the author’s ability to pay publication charges, it undermines authorship equity and has significant implications for defining global research trends, specifically from middle-income countries. Hence, the OA model risks amplifying geographic and institutional disparities in ecology and conservation science, despite its egalitarian philosophy. Incorporating publishing models that account for individual- and institutional-level variations in funding availability, or providing alternative publishing pathways, can help reduce this disparity.

## 1. Introduction

The spirit of science promotes widespread sharing of ideas and findings. Equitable access to scientific knowledge has been recognised as a prerequisite for addressing global challenges and fostering sustainable development (UNESCO, 2025). The United Nations Sustainable Development Goals (UN-SDG, 2025) underscore the importance of knowledge sharing and international collaboration for promoting equality and prosperity. Achieving this requires not only mechanisms for open data and transparent exchange, but also active engagement across the global North and South (Ross-Hellauer et al., 2022).

Historically, scientific journal publishing operated entirely under a subscription-based (SB) model (Solomon & Björk, 2012). Access to published work was locked behind paywalls for readers, while publishing was generally free of charge for authors. In the early 2000s, hybrid journals were introduced, maintaining subscription models while offering authors the option to pay a fee to unlock their individual articles for free public access (Frank et al., 2023). Today, the landscape is shifting rapidly toward full Open Access (OA), accelerated by initiatives such as cOAlition S and its Plan S framework (https://www.coalition-s.org).

The Open-Access (OA) publication model promises to promote diversity, equity, and inclusion in readership by enabling free dissemination of research to anyone, anywhere in the world (Ryan, 2020). Gold OA makes the published article immediately and freely available on the publisher’s website by charging the authors an Article Processing Charge (APCs). In contrast, Green OA relies on author self-archiving – authors publish in conventional subscription journals but upload their manuscripts to institutional or subject repositories for free public access, usually after publisher-imposed embargo periods. The Diamond (or Platinum) OA represents a non-commercial, community-driven model in which articles are immediately free to read and publish, with no direct costs to authors. These may be sponsored by universities or agencies/societies.

Despite several pathways to the OA system, for-profit publication models (Berger, 2021), primarily the Gold Open-Access model, remains the most common. In this model, publishers support publications from developing countries, often by offering APC discounts and waivers based on the first author’s affiliation. For example, authors from countries classified as low-income by the World Bank generally receive a full waiver on APCs, while those classified as lower-middle-income receive a discount. However, the exact list of eligible countries can vary across journals. Therefore, with its egalitarian philosophy, the OA model has been widely appreciated with demonstrated increase in readership (Tang et al., 2017), especially in low- and middle-income countries (Huang et al., 2024). Yet the advantage of free readership for institutions lacking the financial resources to subscribe to journals can be quickly undermined by emerging obstacles to authorship faced by their researchers (Ross-Hellauer et al., 2022). APCs can be extremely high in low- and middle-income countries ($1,500–$3,000 USD per article on average; Frank et al., 2023), even after discounts (Pai, 2020; Solomon & Björk, 2012). In some cases, APCs can even reach up to $10,000 USD, which can be challenging for authors globally. Therefore, although aimed at promoting equity in readership, the OA publication model has been criticised as a discriminatory business model. The high APC can act as a publication barrier for young researchers, especially from resource-poor countries (Frank et al., 2023; Vervoort et al., 2021). Given the potential for resource-based polarisation of publication dynamics and the widening of the digital divide across countries (Ross-Hellauer et al., 2022), the OA publication model can become unsustainable in the long term (Tennant et al., 2016). While such theoretical debates are common, empirical evidence on the impact of the OA model on publication dynamics across countries has been scarce.

Huais et al. (2026) found that journals transitioning to “pay-to-publish” Open Access (OA) show a sharp 46% decline in first-authored papers from non-waiver-eligible countries. However, they acknowledge that their study is largely limited to the publication dynamics of 14 top high-impact journals published by large commercial publishers, and that finer differences across countries eligible for discounts and waivers were not quantified. This is critical as broad trends from select countries may not be representative of global country-level impacts of the OA publication model. Our work directly addresses these gaps by using a massive dataset of 831 journals (1,117,978 publications between 2005 and 2025) to explicitly model publication rates across country-income groups based on both Research4Life and World Bank categories. Additionally, we model these trends across journals with varying APCs and assess how publication charges might directly impact publication dynamics. We hypothesised that *the OA publishing model differentially affects publication dynamics across countries based on their economic strata and journal publication charges*. We predicted that the difference between SB and OA publication rates would be highest in middle-income countries (generally eligible for a discount on the APC), because APCs, even after discounts, would be costly, and that this difference would increase with increasing journal publication charges. Similarly, in journals switching to the OA publication model, publication rates would decrease the most in middle-income countries, followed by high-income countries (not eligible for discounts or waivers), but not in low-income countries (which receive a full waiver on the APC), and these differences would increase with increasing journal APCs.

## 2. Materials and methods

We first downloaded (on the 24^th^ of December 2025) a list of journals in ecology and evolution curated by <u>Research.com</u> (https://research.com/journals-rankings/ecology-and-evolution). The journals on this platform are indexed through a rigorous process based on both their impact and publication volume and so represent a comprehensive list of journals in the field (see https://research.com/our-methodology for more details). We then extracted the source ID, journal number, and open access status (fully Open Access vs hybrid) for all journals indexed in the Scopus database (https://www.scopus.com/sources), and visited journal websites to record a) journal impact factor (as recorded in December 2025), b) journal APC for a full length article (assumed to be 8 pages in length for journals charging on a per page basis), c) whether the journal started as a fully open access journal or transitioned into the model, e) journal start and open-access transition year (if applicable), f) main publisher of the journal. We excluded journals for which APC values were not available or nonexistent (Diamond Access) to obtain the final journal list for our analyses. For details on articles published in these journals, we used a custom Python script (v 3.9) to download the past 20 years of publications (2005-2025) for each journal. From each publication, we extracted details including a) the first author’s institutional affiliation and country, b) publication year, c) publication type (open access vs subscription), and d) unique identifier (DOI).

To explicitly examine whether APC differences correspond to IF differences (thereby providing a benefit of readership to authors) in open-access vs hybrid journals in terms of their impact factor (IF), we assessed differences in APC and IF between open-access and hybrid journals across the two categories for the final journal list using the Wilcoxon rank-sum test.

We sought to understand how a country’s economic status may influence its publication dynamics through eligibility for discounts and waivers on article publication charges, and how these effects may vary across journals with different APC levels. For this, we classified each independent publication into economic strata based on the first author’s institutional affiliation and grouped all selected journals into categories by their APC. Journals generally follow the Research4Life (https://www.research4life.org/access/eligibility/) and/or the World Bank classification (https://blogs.worldbank.org/en/opendata/who-moves-up-and-why--a-closer-look-at-the-new-world-bank-group-) to determine a country’s eligibility for discounts or waivers. Countries in Group A (Research4Life) and/or classified as low income (World Bank) are eligible for automatic waivers (no APC; e.g., Myanmar, Sudan, Sierra Leone), while those in Group B and/or defined as lower-middle income (World Bank) are eligible for discounts (up to 50% of the APC; e.g., Sri Lanka, Viet Nam, Egypt). Upper-middle and high-income countries (World Bank) are grouped together and generally do not receive discounts or waivers (e.g., India, China, USA). We classified countries in our database as high-income (including upper-middle-income), middle-income, and low-income based on the definitions above. The specific list of eligible countries for discounts and waivers may vary across publishers and journals. Therefore, our categorisation only broadly represents country groups, as journals can modify them by adding, removing, or reassigning countries across these categories. Journal APC was incorporated into our analyses by classifying journals into four Tiers based on their APC quartiles calculated across our final list of journals.

We used two complementary modelling approaches to test our predictions (Figure 1). First, for journals maintaining a constant publishing model over the 20-year study window (Non-transitional journals), annual publication counts per journal and country group were modelled as an interaction of country group (low-, middle-, and high-income), journal APC tier (T1, T2, T3 and T4) and paper open-access status (OA vs. SB). Second, for journals transitioning to a fully open-access model (Transitioning journals), annual publication counts were modelled as an interaction of country group (low-, middle-, and high-income), journal APC tier (T1, T2, T3 and T4), and publication timeframe (before vs after the transition). To account for peer-review and publication lags (Forti et al., 2021), articles published up to one year post-transition were categorised as ‘Before OA’, whereas subsequent publications were designated as ‘After OA’. This longitudinal sample was restricted to journals transitioning prior to 2023 to ensure adequate post-transition observation windows. In both cases, we included only publications after 5 years from the journal’s start. This was done because preliminary assessments indicated that publication rates generally saturate after this timeframe, ensuring our dataset omitted the effects of early journal inception and growth.

**Figure 1:**
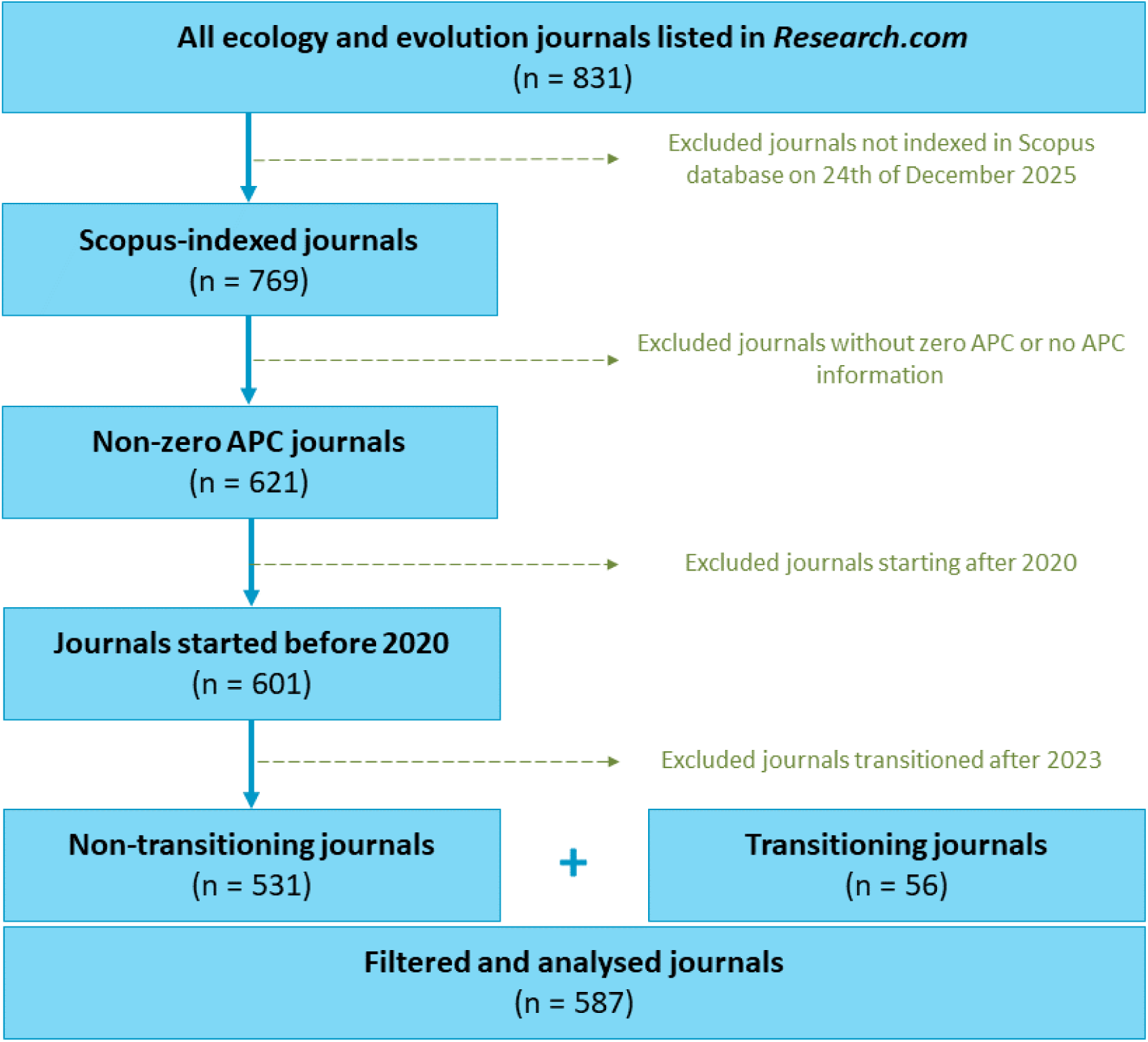
A schematic of the methodology used to filter the Scopus journal dataset and obtain relevant journals for analysis.

In both analytical approaches, we implemented Truncated Poisson Generalised Linear Mixed Models (GLMMs) with log-link functions using the *glmmTMB* package (Brooks et al., 2017) in R (R Core Team, 2025). Models included crossed random intercepts for journal identity and publication year to adjust for journal-specific heterogeneity and temporal trends. Model diagnostics were conducted using the *DHARMa* package (Hartig, 2026) to verify residual uniformity and the absence of overdispersion. In both analyses, we used model parameters and predicted marginal means using the *emmeans* package (Lenth & Piaskowski 2026) to estimate differences across categories and APC tiers, which formed the basis of our inferences.

## 3. Results

We obtained a list of 831 journals in ecology and evolution from Research.com, of which 769 were indexed in Scopus. Our final list included 587 journals (Figure 1), of which 424 were hybrid, 107 were fully open access from their start, and 56 had transitioned to the fully OA model between 2005 and 2023. Cumulatively, our dataset contained 1,117,978 articles across these journals published between 2005 and 2025.

In our dataset, the top 3 publishing houses (Springer, John Wiley & Sons, and Elsevier) owned 50% of all journals. Mean journal APC was $3,006 USD (SD: $1,432). The Wilcoxon rank-sum test suggested that hybrid journals had a significantly greater APC than fully open-access journals (W=54595, p<0.05), but a more variable IF (W=26792, p<0.05; Table 1). The four tiers of APC were T1 (<=$2425), T2 (>$2425 & <=$3190), T3 (>$3190 & <=$3790), and T4 (>$3790). Details of the number of publications in each APC tier and country group are provided in Table S1.

**Table 1:** A summary of Article Publication Charge (APC) and Impact Factor (IF) for fully Open Access and Hybrid journals in our final dataset.

| Journal Type | Mean (SD) APC (USD) | Mean (SD) IF | Count |
| --- | --- | --- | --- |
| Open-Access | 2172 (999) | 2.9 (1.92) | 163 |
| Hybrid | 3326 (1445) | 2.9 (4.49) | 424 |
| Diamond | 0 | 1.12 (0.68) | 117 |

To test for differences between OA and subscription-based publication rates, we fitted a truncated Poisson mixed-effects model with random intercepts for journal and year. The model showed no evidence of overdispersion (DHARMa dispersion test: dispersion = 2.02, p = 0.064), though residual uniformity tests indicated small deviations (Kolmogorov–Smirnov D = 0.11, p < 0.001), as expected in large datasets. Random effects accounted for clustering by journal (variance = 0.94) and year (variance = 0.11). Fixed effects revealed strong influences of country group and open access status with multiple significant interactions (Table S2). Publication rates varied significantly across the three country income groups and four APC tiers (Figure S1). Subscription-based publication rates were, on average, 63% higher than OA publication rates in middle-income countries, 43% in low-income countries, and 38% in high-income countries. Across APC tiers, these differences were lowest in journals in the lowest APC tier (T1, APC <$2425) and either remained stable or increased across APC tiers (Figure 2). Open-access publication rates were higher than subscription-based rates only for low-income countries in T1 APC journals.

**Figure 2.**
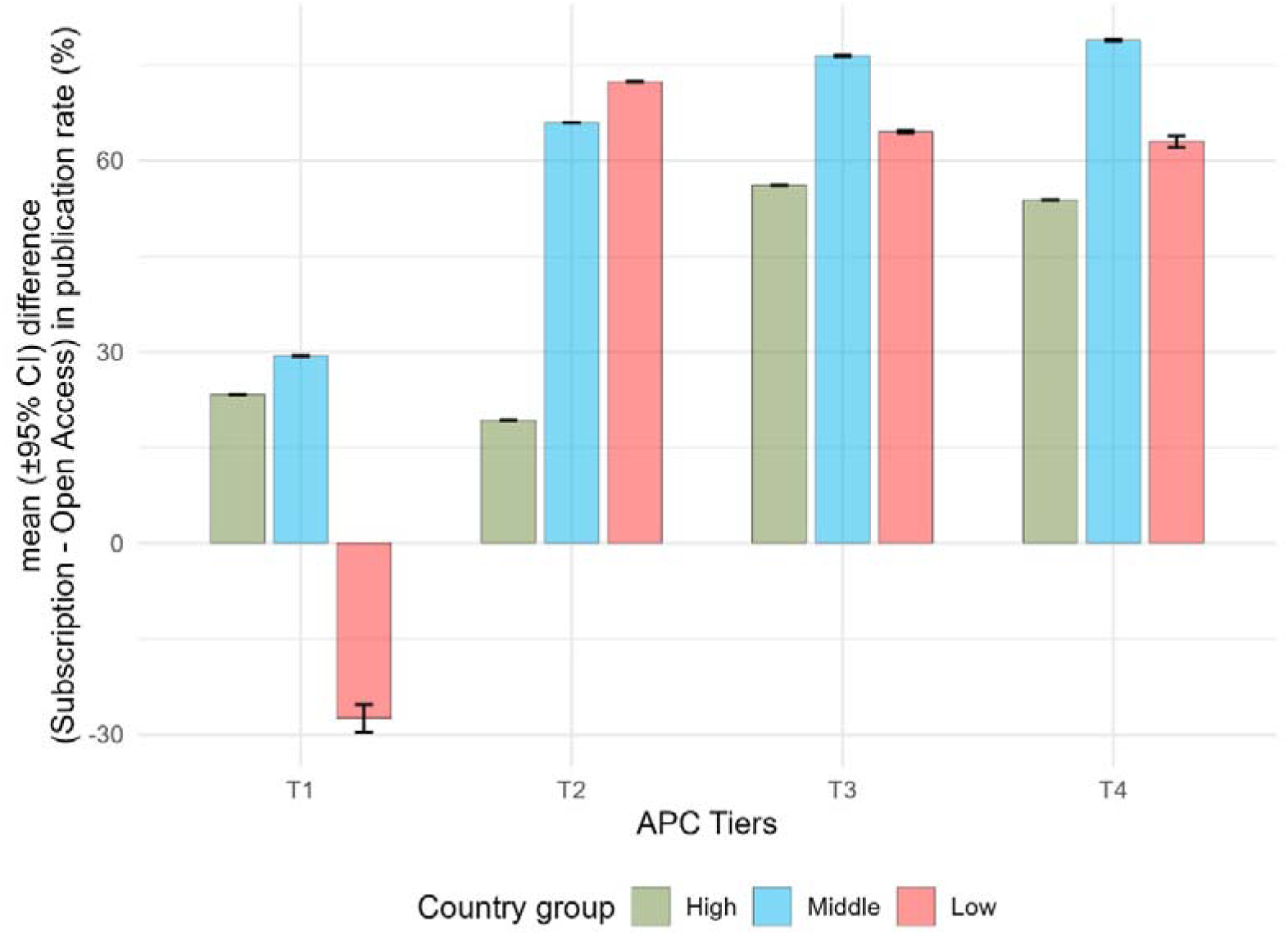
Differences in publication rates for non-transitioning journals. APC tiers are arranged in order of increasing Article Publication Charges.

As above, for publications in transitioning journals, we fitted a truncated Poisson mixed-effects model with random intercepts for journal and year to test for differences between publication rates before and after a journal transitioned into fully open access from a hybrid publication model. The model showed no evidence of overdispersion (DHARMa dispersion test: dispersion = 1.55, p = 0.28), though residual uniformity tests indicated minor deviations (Kolmogorov–Smirnov D = 0.04, p = 0.03). Random effects accounted for clustering by journal (variance = 0.56) and year (variance = 0.39). Fixed effects revealed strong influences of country groups, and publication rates before and after the journal transition varied across country groups (Table S2). On average, publication rates after the journal transition were 19% and 32% lower than before in high- and low-income countries, respectively, but remained unchanged for middle-income countries. Nevertheless, there was significant variation in these trends across APC tiers (Figure 3). For low-APC journals (Tier 1), post-transition publication rates declined across all country groups, most sharply in low-income nations. In higher APC tiers, these declines narrowed, with Tier 3 journals showing post-transition increases led by low-income countries. There was only one journal (with 23 publications) in our list of Tier 4 transitioning journals, which was removed from analysis to ensure model convergence and stability.

**Figure 3:**
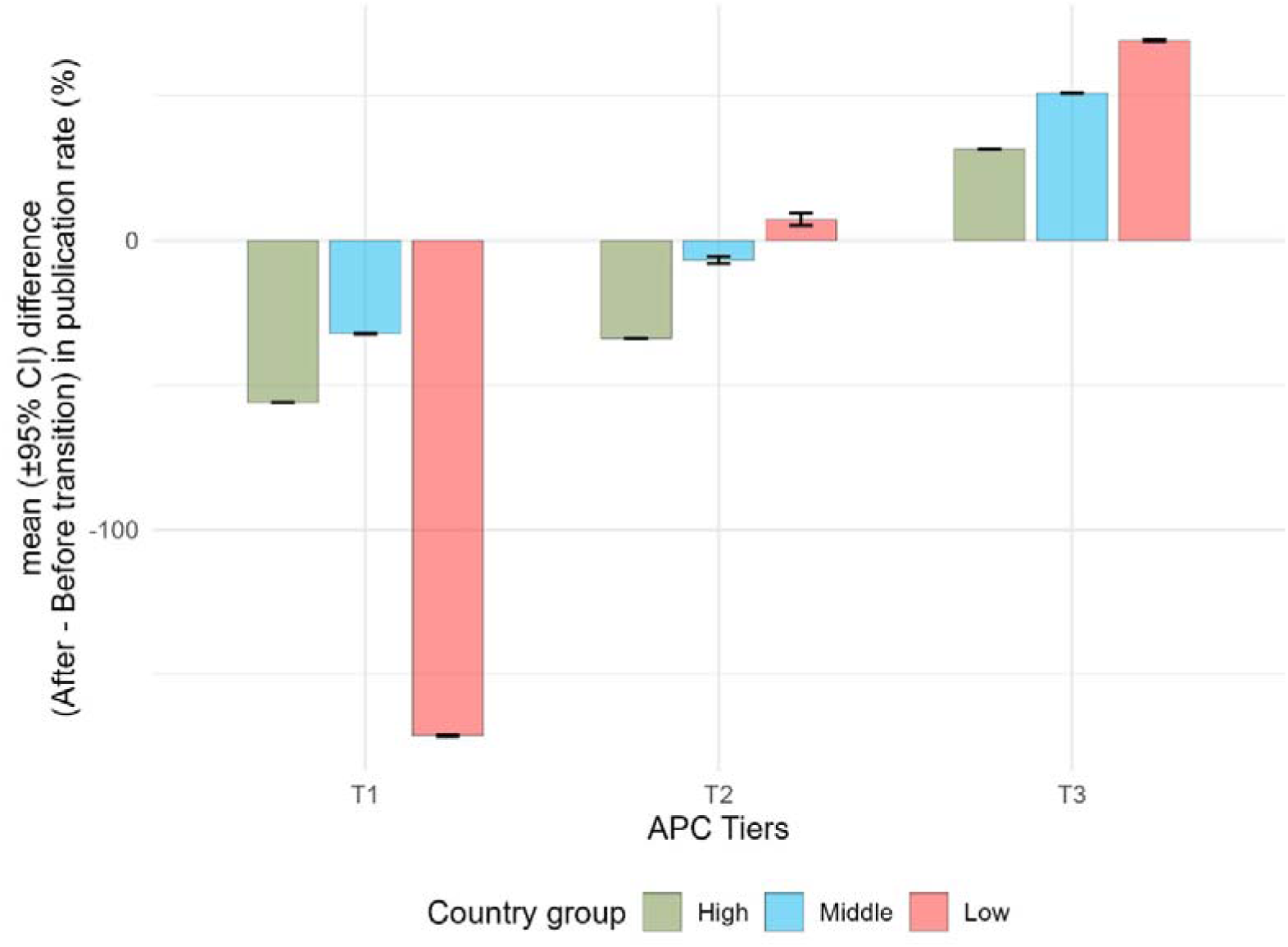
Differences in publication rates for non-transitioning journals. APC tiers are arranged in order of increasing Article Publication Charges.

## 4. Discussion

We analysed over 1.1 million publications over the past 20 years to understand differences in publication rates between subscription and open-access publication models across income-based country groups, and as journal APCs increased. As predicted, we found that subscription-based publication rates were generally higher than open-access publication rates, with the differences greatest in middle-income countries. These differences were also exacerbated across journal APC tiers (i.e., as journal APCs increased). Similarly, for journals that transitioned into the open-access model, overall publication rates decreased after transition. However, in contrast to our predictions, these differences were highest for low-income countries, and publication rates also increased after transition for journals with high APCs (Tier 3). We discuss the implications of our results for open-access publishing of ecological research and suggest potential pathways to more equitable publishing.

### 4.1. Open-Access publication rates are limited by resource availability

We found that for non-transitioning journals, Subscription-Based (SB) publication rates were higher (by 48%) than Open-Access (OA) publication rates across country groups (Figure 2). The growing gap in publication rates between OA and SB models across higher APC tiers (Figure 2) highlights how publication costs strongly influence OA publishing trends. In fact, the rising cost of APC is recognised as an important issue that can offset OA benefits, such as increased readership and citations (Tennant et al., 2016), which OA is promised to provide (Copiello, 2020). It is important to note that we did not find such a trend: the average APC for an open-access journal was $2,172, yet its impact factor (as estimated by Clarivate) was comparable to that of a hybrid journal. The OA option, therefore, seems even less appealing. In fact, journal APCs are driven less by reputation or impact than by the market capitalisation of their publishers (Budzinski et al., 2020). National or institute-level contracts with the publisher, or availability of discretionary or personal funds (Borrego, 2023; Solomon & Björk, 2012), may promote OA publishing, but only for a small proportion of authors with funds.

The trends we observed were more variable among journals that transitioned to the open-access model during our study period. Publication rates decreased significantly across country groups for Tier 1 APC journals (APC < $2,425), but increased for Tier 3 APC journals (APC > $3,790) (Figure 2). This highlights a clear divide: authors with sufficient funding to cover high APCs can maintain or even expand their publication output as journals convert to fully OA models, whereas those with likely constrained budgets cannot. Given the strong correlation between APC and Impact Factor (Spearman’s ρ = 0.62) among transitioning journals, publishing in higher-tier journals likely offers authors a notable readership advantage, albeit at a steep financial cost. Consequently, the sharp decline in publication rates for Tier 1 transitioning journals underscores how the benefits of Open Access remain inequitably tied to institutional and financial resources. Nevertheless, we acknowledge that there could be alternative contributing factors as well, such as journal quality, readership, and publication speed, which may be considered more important than OA (Solomon & Björk, 2012) as long as funding to cover the APC is available, which we could not account for.

### 4.2. Significant impact of OA publishing models on developing countries

The difference in publication rates between OA and SB models was highest for the middle-income country group (63%), followed by the low-income (43%) and high-income (38%) country groups (Figure 2). The large difference suggests that APCs may be prohibitive for researchers, discouraging them from publishing in OA. High-income countries may have greater funding to cover APC, which could explain why these differences were relatively lower for them. A smaller gap in publication rates for low-income countries may indicate the positive impact of APC waivers on publications, but discounts offered to middle-income countries appear to provide less significant relief for authors, who generally lack personal funds and institutional support. In middle-income countries, the APC can equate to several months’ worth of a researcher’s salary even after a discount (Pai, 2020). Indeed, in many such countries, an entire year of field-based research is often funded by modest grants ranging from $5,000 to $10,000 USD, underscoring just how prohibitively expensive APCs can be in comparison.

Huais et al. (2026) observed that post-transition publication rates dropped significantly more in the ‘Global South’ than in the ‘Global North’. While our results align with this finding – since the journals in their dataset correspond to Tier 1 and Tier 2 APCs in our study – we extend their work by evaluating trends across finer income levels (middle- vs. low-income countries). This distinction is vital because these groups receive different financial support (discounts versus full waivers), allowing us to assess the efficacy of each policy and demonstrate how publication disparities vary across higher APC tiers.

While the gap between publication rates for OA and SB is smaller in low-income countries, they contributed only 14.4% of all publications in our dataset, compared to middle-income countries, which produced 28.7%. The actual cumulative research output from the developing world is likely greater, as middle-income countries tend to publish in low-impact and regional journals, often in their native languages (Ekdale et al., 2022). Although important, such journals are rarely indexed by platforms such as Scopus and may even be conflated with predatory journals (Huttner-Koros, 2015), and consequently were out of scope for our analysis. As middle-income countries publish less in OA, they would now be more likely to publish in regional or SB journals, further reducing their discoverability and perceived quality (Ekdale et al., 2022). Therefore, the OA model is likely to result in an overall decrease in publication throughput and availability from the developing world.

Although our analysis examines publication rates at the country level, they might differ across institutions as well. For example, among the top 500 institutions publishing in OA, only 35 (excluding China, which had 66 institutions) were based in middle- or low-income countries. Whereas the top 500 institutions for SB publications featured contributions from at least 53 institutions (excluding China, with 69 institutions) across low- and middle-income countries. Over 50% higher participation from institutes across low- and middle-income countries underscores that the absence of publication charges under the SB model can enable not only higher but also more diverse contributions from the developing world.

The list of countries eligible for waivers varies across journals, and our country-specific analysis cannot directly account for such variation. For ineligible countries according to Research4Life or World Bank classifications, journals may still offer waivers on a case-by-case basis. However, such waivers may be few and difficult to obtain, and more importantly, as our results show, they are insufficient to offset publication bias. In middle-income countries, where the gap in publication dynamics was the largest, even if institutions can offset APC charges with other sources, this is likely to further strain the already small pool of research funds available to them. Nevertheless, only a few institutes may have dedicated publishing funds or agreements with publishing houses for OA, resulting in reduced representation from other institutes.

### 4.3. Towards Inclusive and Sustainable Models of Scientific Publishing

As grant policies mandate publishing only in fully OA journals (Frank et al., 2023), more subscription journals are likely to transition to fully OA. 48 of the 70 OA journals in our dataset transitioned to fully OA in the past decade, and the rate of such transitions is increasing (Borrego, 2023). When comparing publication data before and after journal transitions, our analysis controls for journal-related factors such as publication time and quality, thereby clearly highlighting the nuanced impacts of OA across countries. While full waivers remain an important tool for increasing publication equity, discounts alone are not sufficient. This is problematic, as more journals transition to OA, the options for institutions in middle-income countries to publish their work are constrained. Additionally, the shift in publishing parity towards select institutions in high-income countries can have severe consequences for how *global research* is defined and who gets to define it.

The OA model has significant scope for improvement to truly fulfil its goal of promoting diversity, equity, and inclusion. Since funding availability can vary significantly across institutions and authors (Borrego, 2023), solutions that better account for individual variability should be promoted. For example, research that clearly has no funds for publication in its grants could automatically be eligible for waivers and discounts (e.g., the British Ecological Society offers a fee waiver for such research upon a dedicated application). A framework for basing APC charges on average salary trends for researchers and professors in their respective countries could be considered. Instead of defining eligibility for waivers and discounts solely by a country’s economic status, a more equitable approach might be to assess eligibility at the institutional level (Higino et al., 2025). One may argue that these reforms would negatively affect publishing houses, but the journal’s APC is rarely related to publishing costs (Budzinski et al., 2020). Moreover, lower-cost operations are feasible, particularly given that peer review is typically conducted without compensation (but see Naddaf 2026), and no royalties are offered to the publishing researchers. Such reforms may be challenging to implement without considering the oligopoly and market share of publishing houses (Budzinski et al., 2020). In our dataset, 3 publishers (*Springer*, *John Wiley & Sons*, *Elsevier*) housed 50% of all journals, all of which were established in high-income countries. In several journals, editorial teams also lack representation from the developing world (Berger, 2021), further obscuring the needs of such countries.

Nevertheless, with the realisation of the disparity in the open access publication model, a few other options for OA publications have been introduced recently. For example, the Subscribe to Open (https://subscribetoopencommunity.org/) model allows a consortium of libraries to pool funds to subscribe to various journals, and the publisher makes their content available to everyone once a certain subscriber threshold is reached. Similarly, Elsevier’s Geographical Pricing for Open Access model (https://www.elsevier.support/publishing/answer/geographical-pricing-for-open-access) introduces a separate list of country classification based on income groups and offers discounts on publications where all authors are from a single country group. Society journals that charge less but offer higher-quality peer review (Romesburg, 2016) are not typically considered the first option for publication (Huttner-Koros, 2015) and can be promoted. Several OA journals, for example, Conservation and Society, operate on minimal APCs and are instead supported by donors. Such publication models are underexplored and often overshadowed by the dichotomy between OA and hybrid models. Bypassing traditional commercial journal systems completely, Peer Community In (like PCI Ecology, 2025) offers a non-commercial recommendation platform that evaluates and recommends preprints to ensure transparent, rapid peer review at absolutely no cost to authors or readers. Few regions, such as the South Americas, have already been implementing free peer-review and OA models (or diamond OA models), such as the Scientific Electronic Library Online, where training in publishing practices is provided by non-commercial stakeholders and repositories are hosted by universities or public libraries (Berger, 2021). Lastly, publication models that pay reviewers for their services have recently been explored, highlighting improved review quality and publication times without compromising editorial decision-making (Naddaf, 2026).

Addressing these implications is particularly critical in ecology and evolution as ecological processes operate across scales, often transcending geographic and political boundaries. Synthesising data collected in diverse contexts is essential for developing robust, generalisable insights into ecosystem structure and function. If the OA model disproportionately limits contributions from middle-income countries, large areas of the world – often encompassing biodiversity hotspots and climate-vulnerable ecosystems – risk being underrepresented in the global knowledge pool. Such publication asymmetries reinforce existing geographic and thematic biases in ecological research (e.g., temperate systems studied more extensively than tropical ones (Hirschfeld & Fonseca, 2026)), thereby constraining the generality of ecological theory. For conservation science, which requires integrating ecological evidence into management and policy, the risks are even greater (Alston, 2019; Shanker, 2005). Policies and interventions may be shaped by data from high-income countries, while locally relevant knowledge remains unpublished, inaccessible, or devalued (Fuller et al., 2014; Wilson et al., 2016). This imbalance can distort global narratives about biodiversity loss, climate change, and sustainability, weakening conservation practitioners’ capacity in resource-limited settings to design context-specific solutions.

## Acknowledgements

We would like to thank Prof. K. Shanker, Dr M. Gangal and Dr S. Bhattacharyya for their critical inputs on our initial draft.

## Data accessibility statement

All data and codes used for this research are available at https://doi.org/10.6084/m9.figshare.34046436.

