## Supplementary material for "Middle-income countries disproportionately face the burden of the Open Access publication model": Supplemntary Material

**Supplementary Material S1**

Table S1: Total number of publications for each combination of country group and APC quantile. T4 for transitioning journals had too few publications and was removed from the analysis

| **Journal Group** | **APC Quantile / Range (USD)** | **Discount** | **Normal** | **Waiver** |
| --- | --- | --- | --- | --- |
| **Non-Transitioning Journals** | **T1** (<2425) | 1,084 | 2,323 | 566 |
|  | **T2** (2425 – 3190) | 1,897 | 3,884 | 1,014 |
|  | **T3** (3190 – 3790) | 2,453 | 5,087 | 1,288 |
|  | **T4** (> 3790 to 12290) | 2,425 | 5,073 | 1,253 |
| **Transitioning Journals** | **T1** (< 2425) | 322 | 508 | 186 |
|  | **T2** (2425 – 3190) | 164 | 275 | 95 |
|  | **T3** (3190 – 3790) | 80 | 92 | 32 |
|  | **T4** (> 3790 to 12290) | 7 | 13 | 3 |

Table S2: Fixed-effect estimates (log scale) and standard errors from the two truncated Poisson mixed-effects models. Model 1 examines differences between open-access and subscription journals (paper OA), while Model 2 compares periods before and after the implementation of open access (Time). Both models include random intercepts for journal and year. Significant predictors and interactions highlight strong effects of country group (reference level: High-income), open access status (reference level: No i.e., subscription-based), APC quantiles (reference level: T1), publication time (reference level: After OA transition), and their combinations.

| **Predictor / Interaction** | **Model 1 (Subs vs OA)** | **Model 2 (Before vs After)** |
| --- | --- | --- |
| **Intercept** | 1.36 (0.12) | 0.79 (0.19) |
| **Country group (Normal)** | 2.30 (0.02) | 2.53 (0.03) |
| **Country group (Waiver)** | -1.82 (0.06) | -1.33 (0.09) |
| **Paper OA (Y) / Time (Before)** | -0.35 (0.02) | 0.28 (0.05) |
| **APC (Q2)** | 0.56 (0.13) | -0.11 (0.25) |
| **APC (T3)** | 0.00 (0.12) | 1.32 (0.37) |
| **APC (T4)** | 0.01 (0.13) | — |
| **Country group (Normal) × Paper OA (Y) / Time (Before)** | 0.08 (0.02) | 0.17 (0.05) |
| **Country group (Waiver) × Paper OA (Y) / Time (Before)** | 0.59 (0.07) | 0.72 (0.13) |
| **Country group (Normal) × APC (Q2)** | -0.43 (0.02) | 0.32 (0.09) |
| **Country group (Waiver) × APC (Q2)** | 1.14 (0.07) | 0.93 (0.17) |
| **Country group (Normal) × APC (T3)** | 0.28 (0.02) | 0.39 (0.07) |
| **Country group (Waiver) × APC (T3)** | 0.60 (0.07) | -0.13 (0.17) |
| **Country group (Normal) × APC (T4)** | 0.47 (0.02) | — |
| **Country group (Waiver) × APC (T4)** | 0.41 (0.07) | — |
| **Paper OA (Y) / Time (Before) × APC (Q2)** | -0.73 (0.03) | -0.21 (0.11) |
| **Paper OA (Y) / Time (Before) × APC (T3)** | -1.10 (0.03) | -0.99 (0.09) |
| **Paper OA (Y) / Time (Before) × APC (T4)** | -1.21 (0.04) | — |
| **Country group (Normal) × Paper OA (Y) / Time (Before) × APC (Q2)** | 0.78 (0.03) | 0.06 (0.11) |
| **Country group (Waiver) × Paper OA (Y) / Time (Before) × APC (Q2)** | -0.80 (0.08) | -0.86 (0.23) |
| **Country group (Normal) × Paper OA (Y) / Time (Before) × APC (T3)** | 0.54 (0.03) | 0.17 (0.09) |
| **Country group (Waiver) × Paper OA (Y) / Time (Before) × APC (T3)** | -0.18 (0.09) | -1.18 (0.25) |
| **Country group (Normal) × Paper OA (Y) / Time (Before) × APC (T4)** | 0.70 (0.04) | — |
| **Country group (Waiver) × Paper OA (Y) / Time (Before) × APC (T4)** | -0.03 (0.10) | — |

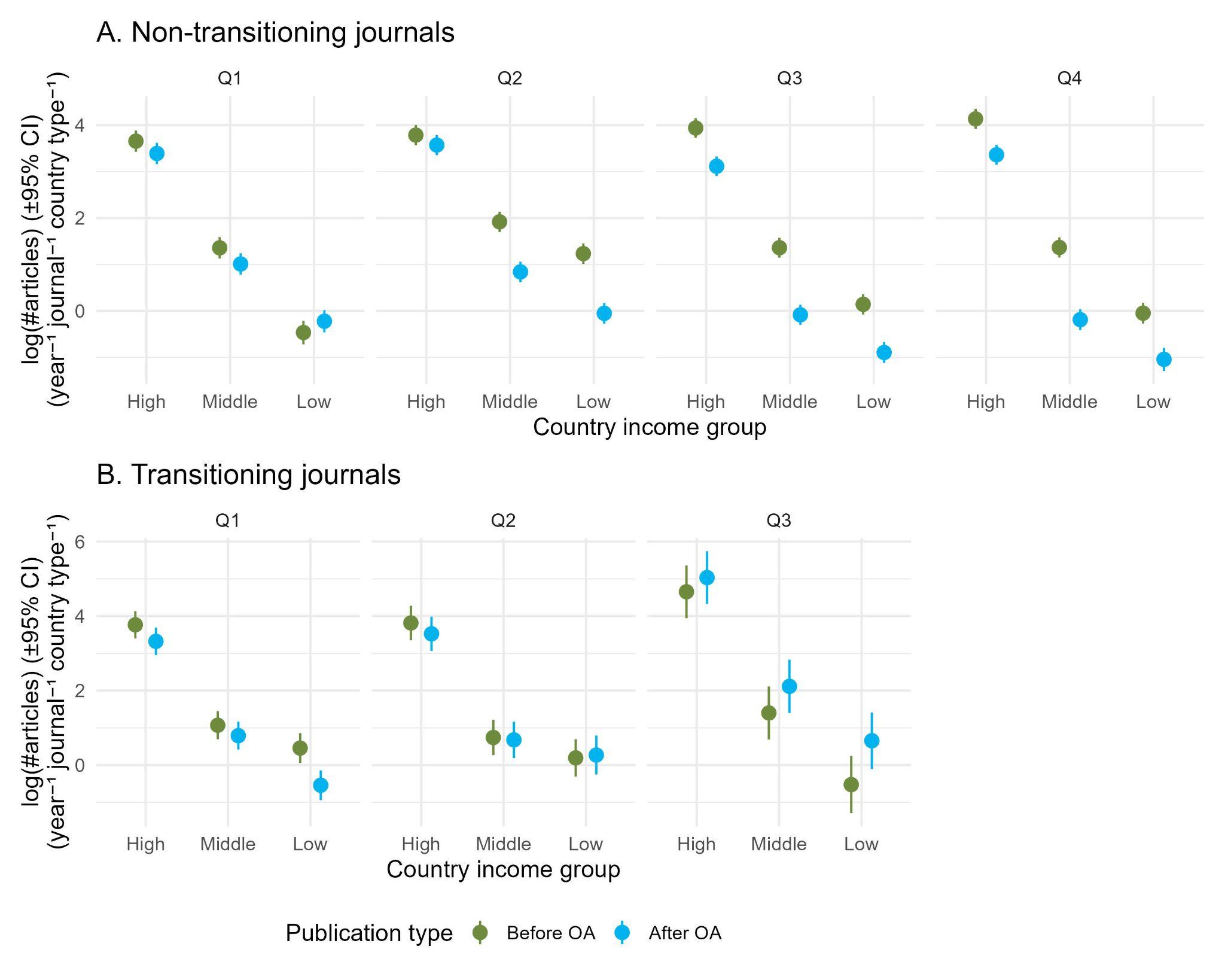

**Figure S1: Publication rates for subscription vs open access publications (left panel) and journals switching to fully open access (right panel). The values in each graph represent marginal means (± 95% CI) for each combination obtained from their respective GLMMs.**
